# Ebselen Exhibits Antimicrobial Activity Against *Clostridioides difficile* By Disrupting Redox Associated Metabolism

**DOI:** 10.1101/2020.07.27.224337

**Authors:** Ravi K.R. Marreddy, Abiola O. Olaitan, Jordan N. May, Min Dong, Julian G. Hurdle

## Abstract

High recurrence rates and spread of antibiotic-resistant strains necessitate the need for alternative therapeutics for *Clostridioides difficile* infections (CDIs). Ebselen, a reactive organoselenium compound inhibits *C. difficile* virulence toxins TcdA and TcdB, by covalently binding to their cysteine protease domains. Ebselen is thought to lack antibacterial activity against *C. difficile* cells and its anti-toxin action is suggested to be solely responsible for its efficacy. However, *C. difficile* has several essential cysteine-containing enzymes that could be potential sites for covalent modification by ebselen; hence, we re-evaluated its anti-*C. difficile* properties. In BHI agar, ebselen inhibited almost all *C. difficile* strains (MICs of 2-8 µg/ml), with ribotype 078 being intrinsically resistant (MIC>64 µg/ml). Wilkins-Chalgren and Brucella agars are recommended media for anaerobic susceptibility testing. Ebselen was either slightly attenuated by pyruvate found in Wilkins-Chalgren agar or obliterated by blood in Brucella agars. Transcriptome analysis showed ebselen altered redox-associated processes, cysteine metabolism and significantly enhanced the expression of Stickland proline metabolism to likely regenerate NAD^+^ from NADH. Intracellularly cells increased the uptake of cysteines, depleted non-protein thiols and disrupted NAD^+^/NADH redox ratio. Growth inhibitory concentrations of ebselen also reduced toxin and spore production. Taken together, ebselen has bactericidal activity against *C. difficile*, with multiple mechanisms of action that negatively impacts toxin production and sporulation. To harness the polypharmacological properties of ebselen, chemical optimization is warranted, especially to obtain derivatives that could be effective in severe CDI, where intestinal bleeding could occur.

## INTRODUCTION

*Clostridioides difficile* is a Gram-positive, spore forming anaerobe and the leading cause of antibiotic associated diarrhea in hospitals (1). *C. difficile* virulence results from production of two glucosyltransferase toxins (Toxin A [TcdA] and Toxin B [TcdB]) (2). These toxins disrupt intestinal epithelial cells by glycosylating Rho/Rac family of GTPases, triggering actin condensation and loss of tight junctions. This leads to inflammation and increased permeability of the gut, tissue necrosis and diarrhea (2). For about 40 years, the antibiotics metronidazole and vancomycin have been the front-line treatments for CDI. These antibiotics being broad-spectrum in action further disrupts the gut microbiota during treatment. This increases the risk of recurrent CDI (rCDI) that occurs in ≥20% of CDI patients (3). Furthermore, infections with epidemic strains, particularly the epidemic B1/NAP1/027 lineage, have drastically reduced the ability to treat CDI (4, 5). Therefore, there is a need for novel CDI therapeutics.

The discovery of alternative CDI therapeutics have focused on blocking the toxin action with toxin binders (e.g. tolevamer) (6), vaccines (7), antibodies such as bezlotoxumab, a monoclonal antibody to TcdB (8) and microbiome-based intervention, such as fecal microbiota transplantation (9, 10). More recently, the organoselenium compound ebselen (2-phenyl-1,2-benzisoselenazol-3(2H)-one) was identified from a high-throughput screen for inhibitors of the cysteine protease domains (CPDs) of *C. difficile* toxins (11). Ebselen directly inhibits both TcdA and TcdB, by forming a covalent bond with the active site cysteine in the toxins CPDs. The CPD is important for toxin action, since their activation by host 1D-myo-inositol hexakisphosphate (IP6) triggers autocleavage and release of the glucosyltransferase domains of TcdA and TcdB. Studies in mice showed ebselen could attenuate CDI caused by the lab strain *C. difficile* 630 (11).

Ebselen is a polypharmacological molecule, which is known as a promising therapeutic for multiple disorders, due to its anti-inflammatory, antioxidative and cryoprotective properties (12-15). Ebselen has strong antimicrobial activities against yeast (16) and selective bacteria, namely Gram-positive bacteria (17-19). It also shows multiple modes of actions against yeast, including inhibiting the proton translocation/ATPase activities (16), targeting glutamate dehydrogenase leading to reactive oxygen species (ROS) (13) and activating DNA damage, thereby altering nuclear proteins (20). In the case of bacteria, ebselen was shown to inhibit thioredoxin reductase enzyme (TrxR) by blocking electron transfer from NADPH to their substrates (21). Thioredoxin (Trx) and glutaredoxin (Grx) systems regulate various critical cellular functions, including ribonucleotide reductase (RNR), peroxiredoxins and methionine sulfoxide reductases (22, 23). Disruption of these cellular functions is often lethal, since the RNR protein is crucial for DNA replication by reducing ribonucleotides to deoxyribonucleotides (22), while peroxiredoxins and methionine sulfoxide reductases protects against oxidative stress (23). Bacteria with both Trx and Grx systems need only one system to counteract ebselen exposure, hence it is more active against species containing only the TrxR system (19, 21). Our review of *C. difficile* genomes revealed they possess the Trx system, appearing to lack Grx. Furthermore, *C. difficile* has multiple proteins with active site cysteines. This prompted us to question whether ebselen could have antimicrobial activity against *C. difficile* cells. Our results show that ebselen is bactericidal against *C. difficile*, but this property is masked by the presence of blood in culture medium. Cellular activity was confirmed from changes to the transcriptome and redox associated metabolites, showing that *C. difficile* readjusts its metabolism to combat ebselen-induced oxidative damage.

## RESULTS

### Ebselen displays antimicrobial activity against *C. difficile*

We first confirmed that ebselen protects Vero epithelial cells from TcdB cytopathy (i.e. cell rounding), in a dose-dependent manner (**Fig. S1**); its EC_50_ was 12.93 μM, somewhat 64-fold lower than reported (11), using protein from List Biological Laboratories, Inc. We next evaluated the MICs of ebselen against 12 strains of *C. difficile*, by the Clinical and Laboratory Standards Institute (CLSI) method using Wilkins-Chalgren agar or Brucella agar (supplemented with hemin (5 mg/L), vitamin K (1 mg/L) and 5% (v/v) defibrinated sheep blood. As described in **Table 1**, ebselen was not active against the test strains in Brucella media (MIC is ≥128 µg/ml). In Wilkins-Chalgren agar ebselen only displayed weak to moderate MICs of 16 – 64 µg/ml; except for a ribotype 002 strain, all other strains were slightly susceptibility to ebselen (**Table 1**). Next we tested ebselen in BHI medium, which revealed it had substantial inhibitory activity against most *C. difficile* strains tested, with MICs ranging from 4 – 8 µg/ml (**Table 1**). Only strains belonging to ribotype 078 were refractory to ebselen (MIC = 32-128 µg/ml, **Table S1**). The difference in MICs between the types of media was further investigated. Wilkins-Chalgren agar contains hemin and sodium pyruvate, which we reasoned could alter biological redox processes in *C. difficile*. Hence, we supplemented BHI agars with hemin or pyruvate components. Supplementation with hemin (5 μg/ml) did not alter MICs (*data not shown*), whereas addition of 1 g/L of pyruvate worsened MICs by 2-16-fold (**Table 1**). We further tested the effect of blood used in Brucella agars; the activity of ebselen was ablated (MIC >128 µg/ml; **Table 1**) when BHI agar was supplemented with 5% (v/v) defibrinated sheep blood. In contrast, vancomycin strongly inhibited the growth of all tested strains irrespective of media composition with MICs of 0.5 – 2 µg/ml (**Tables 1** and **S1**). These observations reveal that the activity of ebselen against *C. difficile* is strongly influenced by the composition of the medium used in susceptibility testing and implies environmental conditions could affect its antibacterial activity.

**Table 1.** Antimicrobial activity of ebselen (EBS) and vancomycin (VAN) against various *C. difficile* strains.

| Strain | PCR-ribotype | Agar MIC (µg/ml)<sup>A</sup> |  |  |  |  |  |  |  |  |  |
| --- | --- | --- | --- | --- | --- | --- | --- | --- | --- | --- | --- |
|  |  | BHI |  | BHI + blood |  | BHI + 1g/L<br>pyruvate |  | Brucella |  | Wilkins-Chalgren |  |
|  |  | EBS | VAN | EBS | VAN | EBS | VAN | EBS | VAN | EBS | VAN |
| R20291 | 027 | 4 | 0.5-1 | >128 | 1-2 | 8 | 1 | 128 | 0.5-1 | 32 | 0.5-1 |
| NR49292 | 001-072 | 4 | 0.5-1 | >128 | 1-2 | 16 | 1 | >128 | 1-2 | 64 | 0.5-1 |
| NR49305 | 002 | 8 | 1 | >128 | 1-2 | 16 | 1 | >128 | 1-2 | 128 | 1-2 |
| NR49294 | 014 | 4-8 | 0.25-0.5 | >128 | 2-4 | 16 | 0.5 | >128 | 2 | 32-64 | 0.25-0.5 |
| NR49312 | 017 | 4 | 0.25 | >128 | 1-2 | 32 | 1 | >128 | 2 | 16 | 0.5 |
| NR49323 | 018 | 4 | 1-2 | >128 | 1-2 | 16 | 1 | >128 | 2 | 32-64 | 0.5-1 |
| NR49277 | 019 | 8 | 1-2 | >128 | 1-2 | 16 | 1 | >128 | 2 | 64 | 1 |
| NR49300 | 020 | 4 | 0.5 | >128 | 1-2 | 16 | 0.5 | >128 | 1 | 32-64 | 0.5 |
| NR49317 | 024 | 2-4 | 0.5 | >128 | 1-2 | 32 | 1 | >128 | 1-2 | 32-64 | 0.5 |
| NR49314 | 047 | 4 | 0.25 | >128 | 1-2 | 16-32 | 1 | >128 | 1-2 | 16-32 | 0.5 |
| NR49325 | 054 | 4 | 0.5 | >128 | 1-2 | 16 | 1 | >128 | 2 | 32-64 | 0.25 |
| NR49318 | 106 | 8-16 | 0.5 | >128 | 1-2 | 32 | 1 | >128 | 2 | 32-64 | 0.5 |

<sup>A</sup>MICs are from three biological replicates and shown as the range, where obtained; BHI = brain heart infusion.

### Ebselen is bactericidal

At its MBC of 16 µg/ml ebselen killed >99.9% of logarithmic cells of R20291; thus, ebselen was bactericidal at 8 x its MIC.

### Activity of ebselen against representative gut flora

We tested ebselen against various gut bacterial anaerobes (**Table 2**). It did not primarily inhibit growth of *Bacteroides spp*. and *Porphyromonas uenonis* (MIC is ≥128 µg/ml), but it inhibited the growth of *A. viscosus* (MIC=2 µg/ml), *L. crispatus* (MIC=32 µg/ml), *L. johnsonii* (MICs=8-16 µg/ml), *F. nucleatum* (MICs=4 µg/ml) and *F. periodonticum* (MIC=8 µg/ml). Similar to the observations for *C. difficile*, addition of 5% (v/v) defibrinated sheep blood diminished the activity of ebselen. As a control, vancomycin inhibited these strains with MICs of 0.5 – 8.0 µg/ml, except against *Bacteroides sp*. HM19 (MIC >32 µg/ml). These observations confirm that blood impairs the antimicrobial activity of ebselen and that ebselen may not inhibit growth of *Bacteroides* species.

**Table 2.** Antimicrobial activity of ebselen and vancomycin against a panel of gut bacterial species.

| Bacteria | Strain | MIC (µg/ml)<sup>A</sup> |  |  |  |
| --- | --- | --- | --- | --- | --- |
|  |  | BHI |  | BHI + blood |  |
|  |  | EBS | VAN | EBS | VAN |
| *Actinomyces viscosus* | HM238 | 2 | 0.5 | 64 | 0.5 |
| *Bacteroides fragilis* | HM20 | >128 | 2 | >128 | 8 |
| *Bacteroides ovatus* | HM222 | >128 | 1 | >128 | 1 |
| *Bacteroides sp.* | HM18 | 128 | 4 | >128 | 8 |
| *Bacteroides sp.* | HM19 | 128 | >32 | 128 | >32 |
| *Bacteroides sp.* | HM23 | 128 | 2 | >128 | 4 |
| *Bacteroides sp.* | HM28 | 64 – 128 | 2 | 128 | 32 |
| *Lactobacillus crispatus* | HM421 | 32 | <0.25 | >128 | <0.25 |
| *Fusobacterium nucleatum* | HM260 | 4 | 8 | 64 | 32 |
| *Fusobacterium periodonticum* | HM41 | 8 | 8 | 128 | 32 |
| *Lactobacillus johnsonii* | HM643 | 8 – 16 | 0.5 | 128 | 0.5 |
| *Porphyromonas uenonis* | HM130 | >128 | 1 | >128 | 0.5 |

<sup>A</sup>MICs are from three biological replicates and shown as the range, where obtained; BHI = brain heart infusion.

### Transcriptome analysis of ebselen mode of action against *C. difficile*

Ebselen inhibits bacterial TrxR and also disrupts multiple macromolecular processes in *S. aureus* i.e. DNA, RNA, proteins, lipid and cell wall biosynthesis (17, 19). To better understand how ebselen inhibits *C. difficile* growth, we analyzed the global transcriptional response of *C. difficile* R20291 by RNA sequencing (RNAseq). RNA samples were prepared from three independent biological replicates. The raw data of the analysis is deposited in NCBI database under accession number PRJNA647225. The volcano plot in **Fig. 1A** shows differences in gene expression between the untreated and ebselen-treated cultures. A total of 565 genes (**Fig. 1B** and **Table S2**) were differentially expressed by ebselen exposure (FDR <0.01 and fold change >2.0), of which 360 were upregulated and 205 were downregulated. Transcription of several metabolic pathways were altered, suggesting ebselen imposed a metabolic burden (**Table S2**). Various ribosomal proteins and proteins involved in amino acid metabolism were upregulated. Transcript levels for molecular chaperones were unchanged, suggesting ebselen did not impose a burden on protein misfolding in cells. Proteins involved in DNA mismatch repair and homologous recombination were also upregulated. Ebselen also enhanced transcription of various transporters involved in the uptake of sugars, iron, sulfur and amino acids. Expression levels for flagellar assembly and bacterial chemotaxis were also upregulated. The most remarkable changes were those associated with bacterial redox, as discussed in the below sections.

**Figure 1.**
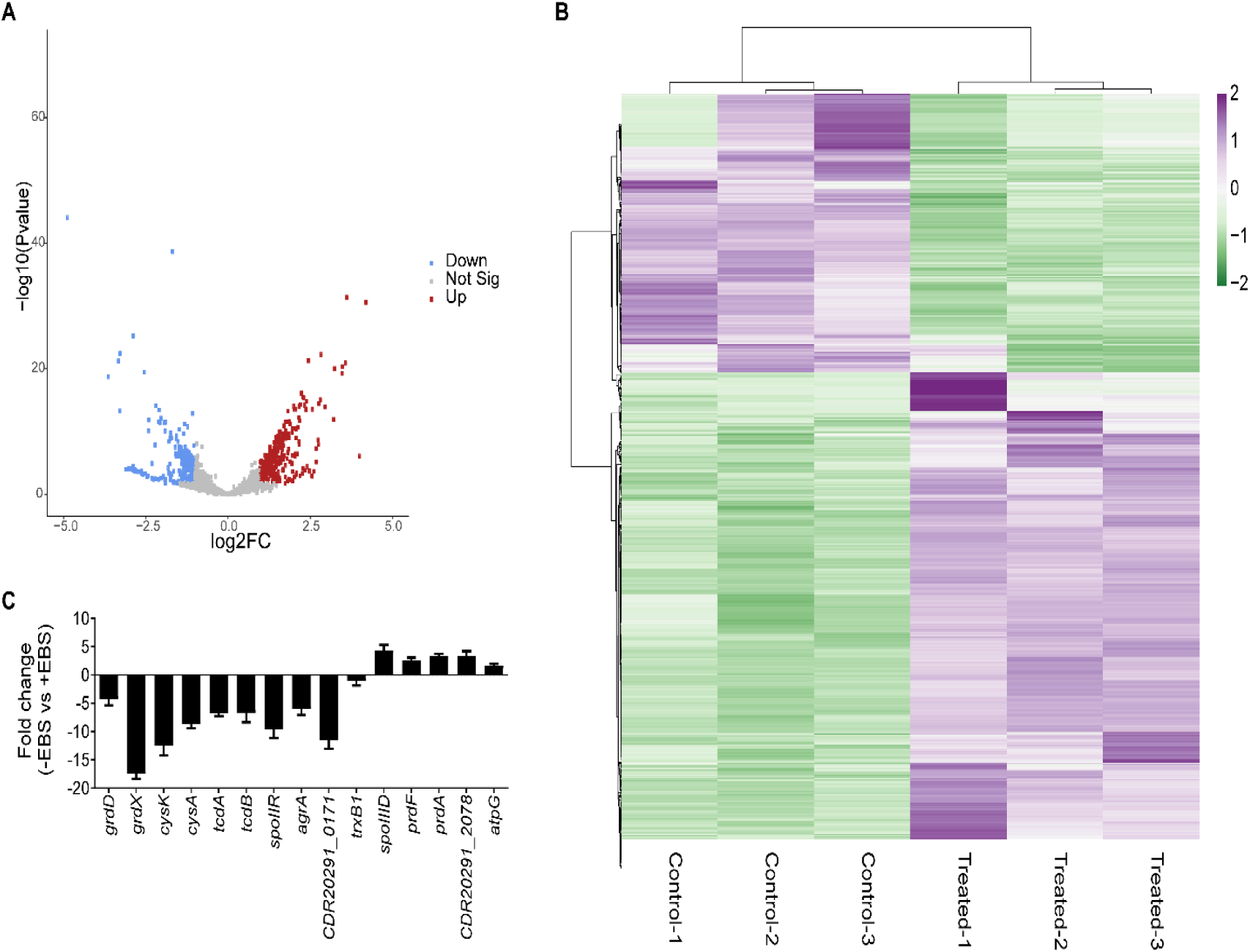
Analysis of global gene expression in presence of ebselen. *C. difficile* R20291 was grown to early exponential phase (OD_600_ ≈ 0.2) and exposed to 16 µg/ml of ebselen for 30 min before RNA was extracted for sequencing. Controls were treated with DMSO. RNAseq data was analyzed on the Galaxy web-based platform. **(A)**The quality of the RNAseq data was analyzed by principle component analysis and data visualized in volcano plots of statistical significance versus fold change. **(B)** Heat map of differentially expressed genes is shown; the color intensity provides a measure of gene expression (purple for upregulated and green for downregulated genes). The heat map was generated using Clustvis software. **(C)** mRNA levels were analyzed for various genes by RT-qPCR, the fold change was calculated as the difference in mRNA levels of control vs ebselen treated cells.

### Ebselen increases cysteine pool, while depleting non-protein thiols

Since *C. difficile* has three TrxR, we hypothesized ebselen might inhibit one or more of these enzymes and alter the expression of thioredoxins (*trx*) genes that depend on TrxR. However, it did not substantially affect the transcription of thioredoxins in the RNAseq (**Table S2 and Fig. 1C**); although, thioredoxin genes were not induced by ebselen, they are known to respond to reactive stress from metronidazole (24). Furthermore, the MICs of ebselen were not shifted by expression of three *trx* genes (trxA1, *trxB1* and *trxB3*) on plasmid pRPF185 in R20291 (*data not shown*). Transcription of genes regulated by the Trx system were also unaffected i.e. expression of RNR, peroxiredoxin (*bcp*) and methionine sulfoxide reductase (*msrAB*) were unaffected. Transcription of superoxide dismutase (*sodA*) was also unaffected suggesting that ebselen did not induce the formation of ROS. Therefore, the anti-*C. difficile* activity of ebselen might be independent of the Trx system. Albeit, biophysical and biochemical characterization will be required to better determine if ebselen inhibits one or more of *C. difficile* TrxRs.

Interestingly, our RNAseq showed that transcript levels for cysteine metabolism was affected, as genes *cysM* (O-acetylserine sulfhydrylase) and *cysA* (serine acetyltransferase) were downregulated by 1.72 and 1.84-fold respectively (**Table S2**). RT-qPCR confirmed that *cysA* was downregulated by 8.7 ± 1.6 – fold (**Fig. 1C**); we also found that *cysK*, encoding another O-acetylserine sulfhydrylase, was downregulated by 12.4 ± 3.99-fold. CysM, CysA and CysK are thought to be involved in cysteine degradation or cysteine biosynthesis from serine (25). Interestingly, expression of the ABC cystine/cysteine transporter subunit (*CDR20291_2078*) was upregulated by 3.96-fold (**Table 3**), which was confirmed by RT-qPCR (3.4 ± 1.6–fold upregulation) (**Fig. 1C**). Based on these findings, we hypothesized that ebselen exposed cells might accumulate cystine/cysteine. We therefore exposed cells to various concentrations of ebselen for 1 h and quantified intracellular concentrations of cysteine. Intracellular cysteine levels were not significantly affected by 2 and 4 µg/ml of ebselen, but 8 and 16 µg/ml enhanced cysteine pools by 5.84 ± 2.13 (p = 0.02) and 7.64 ± 2.1 – fold (p = 0.0036), respectively (**Fig. 2A**). This supports the above hypothesis that cells accumulated cysteine in response to bactericidal concentrations of ebselen. As a negative control, intracellular cysteine levels were not affected by an inhibitory concentration of vancomycin (1 µg/ml).

**Table 3.**
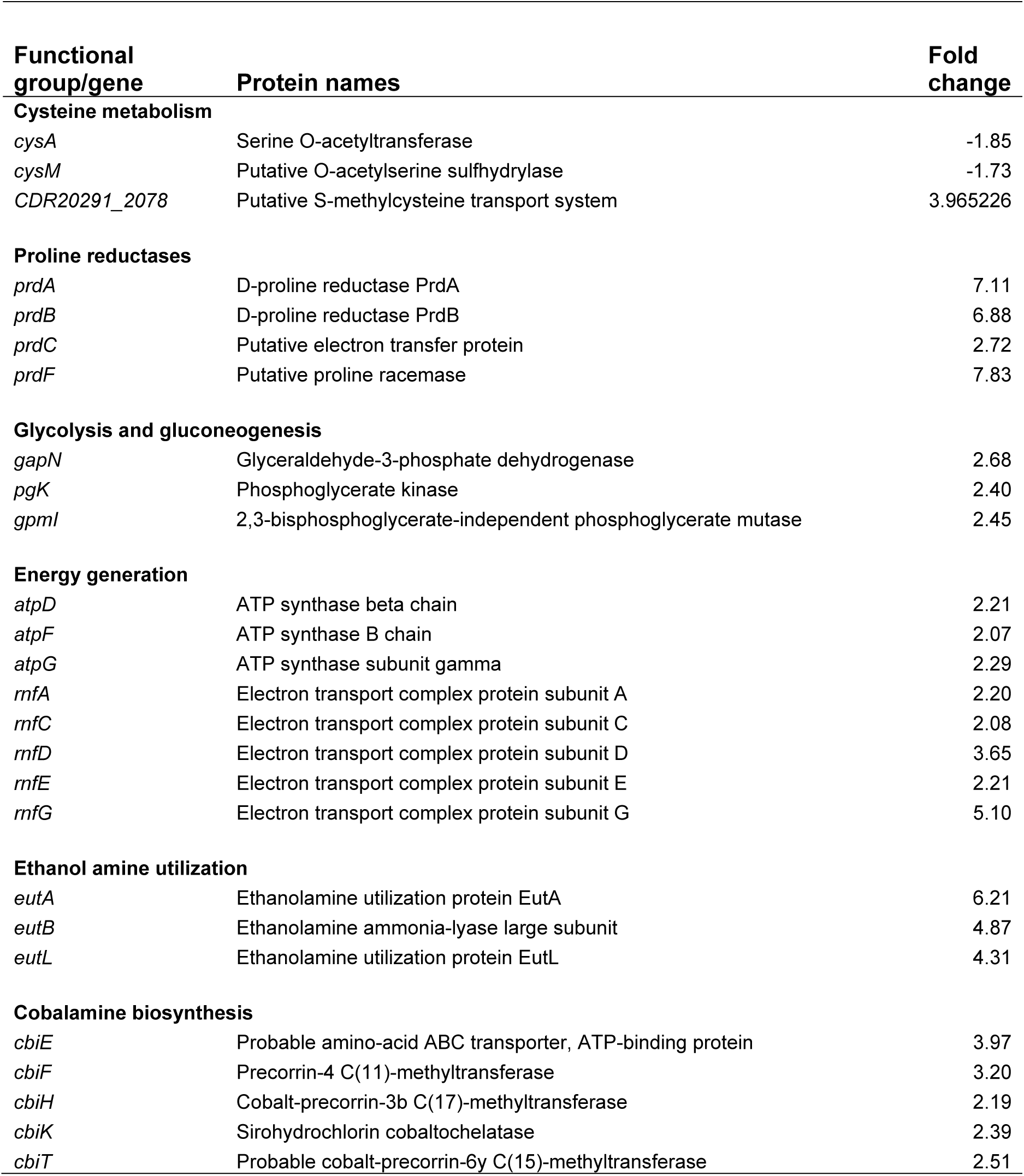
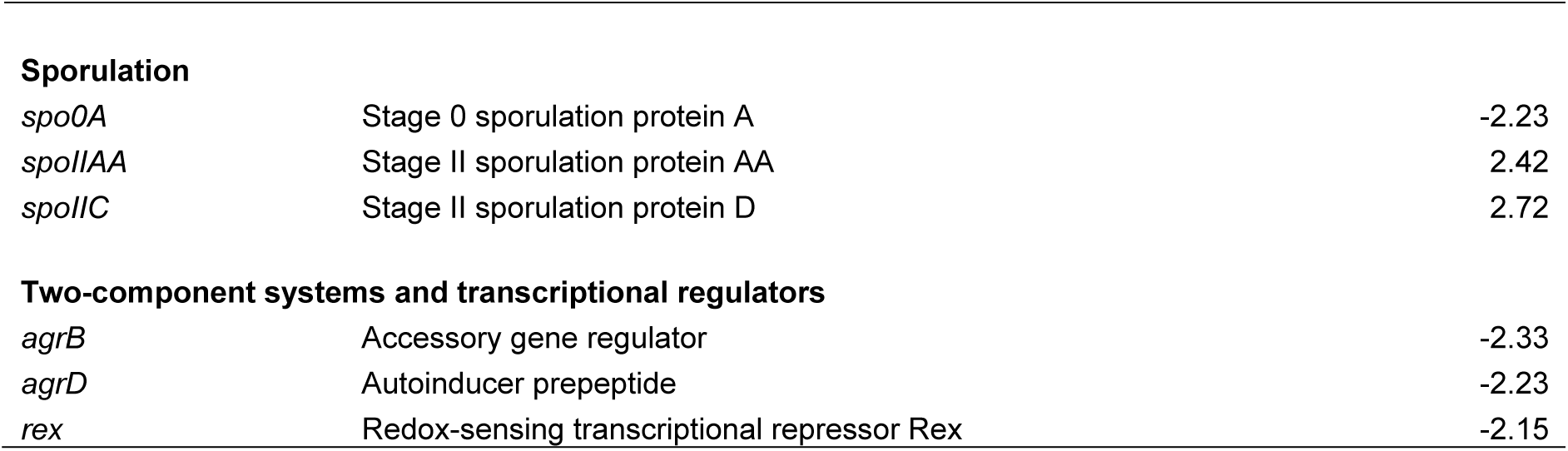
List of selected genes in *C. difficile* R20291 that are differentially expressed by ebselen; their functional classifications are shown.

| Functional group/gene | Protein names | Fold change |
| --- | --- | --- |
| **Cysteine metabolism** |  |  |
| *cysA* | Serine O-acetyltransferase | -1.85 |
| *cysM* | Putative O-acetylserine sulfhydrylase | -1.73 |
| *CDR20291_2078* | Putative S-methylcysteine transport system | 3.965226 |
| **Proline reductases** |  |  |
| *prdA* | D-proline reductase PrdA | 7.11 |
| *prdB* | D-proline reductase PrdB | 6.88 |
| *prdC* | Putative electron transfer protein | 2.72 |
| *prdF* | Putative proline racemase | 7.83 |
| **Glycolysis and gluconeogenesis** |  |  |
| *gapN* | Glyceraldehyde-3-phosphate dehydrogenase | 2.68 |
| *pgK* | Phosphoglycerate kinase | 2.40 |
| *gpmI* | 2,3-bisphosphoglycerate-independent phosphoglycerate mutase | 2.45 |
| **Energy generation** |  |  |
| *atpD* | ATP synthase beta chain | 2.21 |
| *atpF* | ATP synthase B chain | 2.07 |
| *atpG* | ATP synthase subunit gamma | 2.29 |
| *rnfA* | Electron transport complex protein subunit A | 2.20 |
| *rnfC* | Electron transport complex protein subunit C | 2.08 |
| *rnfD* | Electron transport complex protein subunit D | 3.65 |
| *rnfE* | Electron transport complex protein subunit E | 2.21 |
| *rnfG* | Electron transport complex protein subunit G | 5.10 |
| **Ethanol amine utilization** |  |  |
| *eutA* | Ethanolamine utilization protein EutA | 6.21 |
| *eutB* | Ethanolamine ammonia-lyase large subunit | 4.87 |
| *eutL* | Ethanolamine utilization protein EutL | 4.31 |
| **Cobalamine biosynthesis** |  |  |
| *cbiE* | Probable amino-acid ABC transporter, ATP-binding protein | 3.97 |
| *cbiF* | Precorrin-4 C(11)-methyltransferase | 3.20 |
| *cbiH* | Cobalt-precorrin-3b C(17)-methyltransferase | 2.19 |
| *cbiK* | Sirohydrochlorin cobaltochelataase | 2.39 |
| *cbiT* | Probable cobalt-precorrin-6y C(15)-methyltransferase | 2.51 |
| **Sporulation** |  |  |
| *spo0A* | Stage 0 sporulation protein A | -2.23 |
| *spoIIAA* | Stage II sporulation protein AA | 2.42 |
| *spoIIC* | Stage II sporulation protein D | 2.72 |
| **Two-component systems and transcriptional regulators** |  |  |
| *agrB* | Accessory gene regulator | -2.33 |
| *agrD* | Autoinducer prepeptide | -2.23 |
| *rex* | Redox-sensing transcriptional repressor Rex | -2.15 |

**Figure 2.**
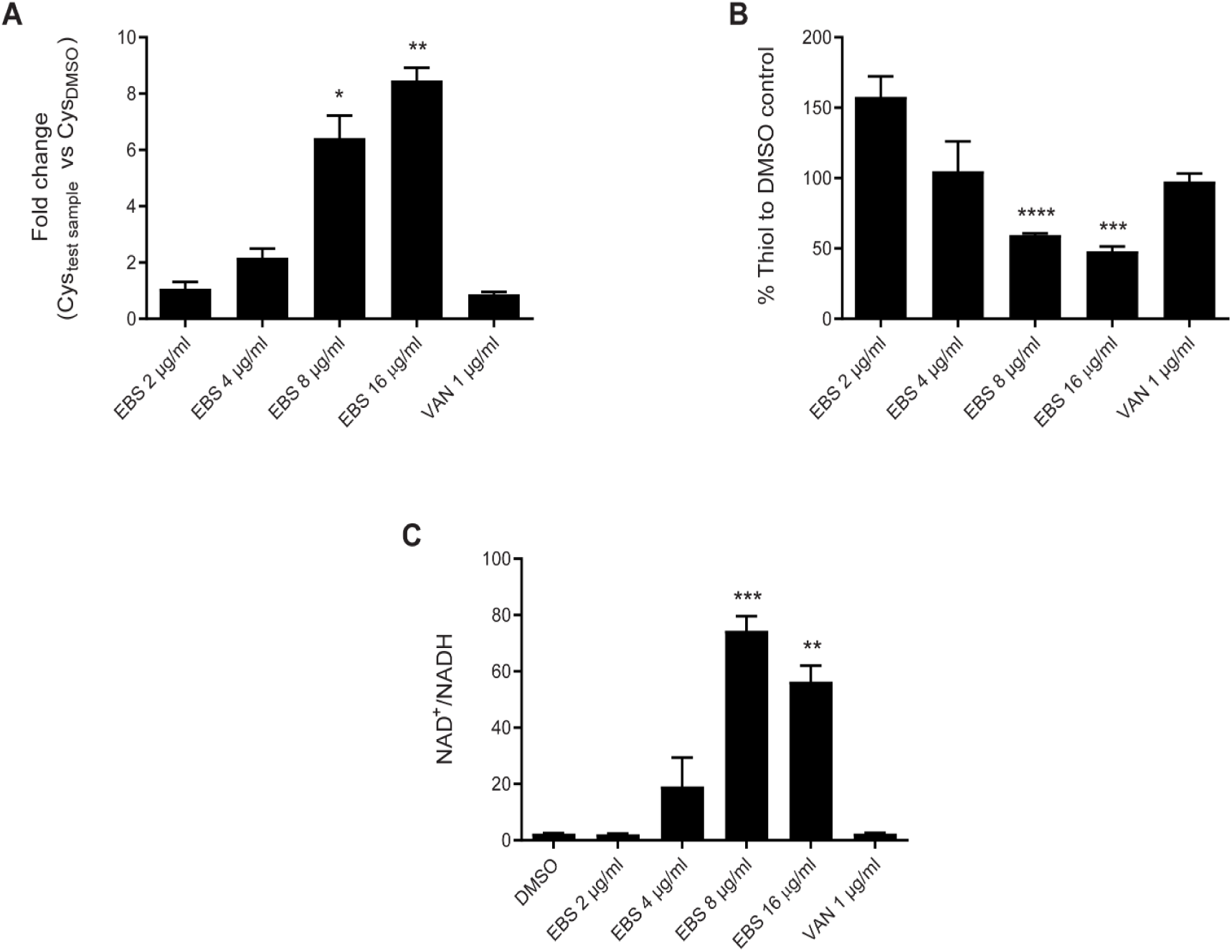
Change in cytosolic content of free cysteine, thiols and NAD^+^/ NADH in presence of ebselen. *C. difficile* R20291 were grown to early exponential phase (OD_600_ ≈ 0.4) and treated with 2, 4, 8 or 16 µg/ml of ebselen. DMSO and vancomycin (1 µg/ml) were used as controls. Whole cell lysates from the same cultures were used to analyze **(A)** cysteine **(B)** protein free thiols and **(C)** NAD^+^/NADH using respective kits from various manufacturers. For cysteine and thiol quantifications the fold change/percent fold change were calculated for the respective test samples relative to DMSO controls. The data represented were normalized to 1 mg of cellular protein content. Error bar indicate means ± SEM; n = 3 (unpaired t-test with Welch’s correction *P<0.05, **P<0.01 and ***P<0.001; done using Graphpad prism version 8.4).

Low molecular weight (LMW) non-protein thiols serve as a defense mechanism against reactive species. To evaluate if ebselen also interfered with non-protein thiols, we quantified the free thiol content of cells. The levels of LMW non-protein thiols were not altered by 2 and 4 µg/ml of ebselen, whereas the thiol content was significantly reduced by 59.6 ± 2.45 (p = <0.0001) and 48.1 ± 6.6 – fold (p = 0.0006) with 8 and 16 µg/ml of ebselen, respectively (**Fig. 2B**). The control vancomycin (1 µg/ml) did not impact the LMW thiol pool (**Fig. 2B**). Based on these observations, we speculate that *C. difficile* adopts LMW thiols to detoxify ebselen and it might transport cystine/cysteine into cells to replete thiol pools.

### Regulation of NAD^+^ generating pathways and energy generation

The most significantly upregulated genes by ebselen belonged to the proline reductase subunit (*prdA, prdB, prdC* and *prdF*), which were upregulated by 3 to 8-fold (**Table 3**). Analysis by RT-qPCR revealed that *prdA* and *prdF* were upregulated by 3.3 ± 1.1 and 2.5 ± 1.2 – fold, respectively (**Fig. 1D**). *C. difficile* proline reductase is a selenoenzyme that is crucial for maintaining cellular redox balance and generating energy via the Stickland pathway (26). The Stickland reaction involves a series of coupled oxidation and reduction of amino acid pairs, in which oxidation of amino acids lead to formation of ATP and NADH while the reductive pathway regenerates NAD^+^ from NADH. Similar to proline reductases, glycine reductase also regenerates NAD^+^ from NADH in the alternate branch of the Stickland pathway (27). RT-qPCR reveled that mRNA for glycine reductase subunits *grdD* and *grdX* were down regulated by 4.2 ± 1.99 and 17.4 ± 2 – fold respectively (**Fig. 1C**), even though these genes were not significantly downregulated in the RNAseq (**Table S2**). Upregulation of proline reductase genes, and downregulation of glycine reductase genes, suggest activation of proline metabolism for the purpose of generating NAD+. Proline reductase is thought to be the preferred route for NAD^+^ regeneration in *C. difficile* (27). We therefore quantified how ebselen influenced the cellular ratio of NAD^+^/NADH. As shown in **Fig. 2C**, ebselen decreased cellular amounts of NADH leading to disruption of the intracellular NAD+/NADH ratio, whereas vancomycin did not have an impact. Proline reduction is thought to also result in a proton motive force by coordinating with the electron transport Rnf complex and the resulting ion gradient may be used to generate ATP by the ATP synthase (28). Consistent with upregulation of proline reductase genes, ATP synthase subunits *atpD, atpF* and *atpG* were also upregulated, along with the electron transport protein complex encoded by *rnfA, rnfC, rnfD, rnfE and rnfG* (**Table 3**). RT-qPCR showed that transcript levels for *atpG* were upregulated by 1.62 ± 0.6-fold (**Fig. 1D**). Hence, in ebselen exposed cells proline may be used for both energy production and NAD^+^ formation.

Other pathways associated with energy metabolism were also altered. For e.g. R20291 exposed to ebselen upregulated genes involved in synthesizing acetyl CoA e.g. genes for ethanolamine utilization (*eutA, eutB* and *eutL*). Conversion of ethanolamine to acetyl CoA requires coenzyme B12 (cobalamin), which may explain the significant upregulation of cobalamin biosynthesis genes (*cbiE, cbiF, cbiH, cbiK* and *cbiT*). This further suggests ebselen altered energy metabolism.

**Table 1** shows that supplementation of BHI with 1 g/L of pyruvate weakened the activity of ebselen activity against *C. difficile*. Pyruvate enters various metabolic pathways in *C. difficile*; for example, it can be converted to acetyl CoA by pyruvate-ferredoxin oxidoreductase (PFOR) encoded by *nifJ*, which was upregulated by 1.69-fold. Alternately, PFOR can convert acetyl CoA to pyruvate, which can be converted to formate by pyruvate lyase, which goes on to generate protons. Unfortunately, the transcriptional snapshot of the cell, and test conditions, did not provide an adequate insight to how pyruvate affects the action of ebselen.

### Growth inhibitory concentrations of ebselen inhibits *C. difficile* toxin production

Various factors are known to influence the expression of *tcdA* and *tcdB* (29). For example, activation of PrdR represses toxin gene expression (30), whereas overexpression of Agr, a two-component regulatory system, enhances the toxin gene expression (31). The *agr* loci comprises of *agrD* that produces a signal peptide, which is recognized by the membrane bound kinase AgrC. Once activated AgrC phosphorylates the DNA-binding response regulator AgrA leading to its dimerization and binding to promoters, thus upregulating the *agr* operon and virulent genes. The Agr operon (*agrB* and *agrD*, 2.33 and 2.23-fold decrease respectively) was downregulated by ebselen (**Table S2**); this was confirmed by RT-qPCR showing *agrA* was decreased by 5.9 ± 2.4-fold (**Fig. 1C**). These observations implied that ebselen could reduce toxin production through PrdR activation and Agr downregulation. RT-qPCR showed that *tcdA* and *tcdB* were downregulated by 6.74 ± 1.14 and 6.68 ± 3.76-fold, respectively (**Fig. 1C**). We next quantified toxins by ELISA using stationary phase cultures that were exposed to a sub-cidal concentration of ebselen (2 µg/ml) for 24 h. As shown in **Fig. 3A**, ebselen (2 µg/ml) inhibited TcdA and TcdB production by ∼50 and ∼40% respectively. At growth inhibitory concentrations (4 and 8 µg/ml), production of both TcdA and TcdB were inhibited by >80% (p < 0.0002). As a positive control, glucose (1% w/v) strongly inhibited toxin production (**Fig. 3A**), in contrast to the negative control vancomycin (1 µg/ml).

**Figure 3.**
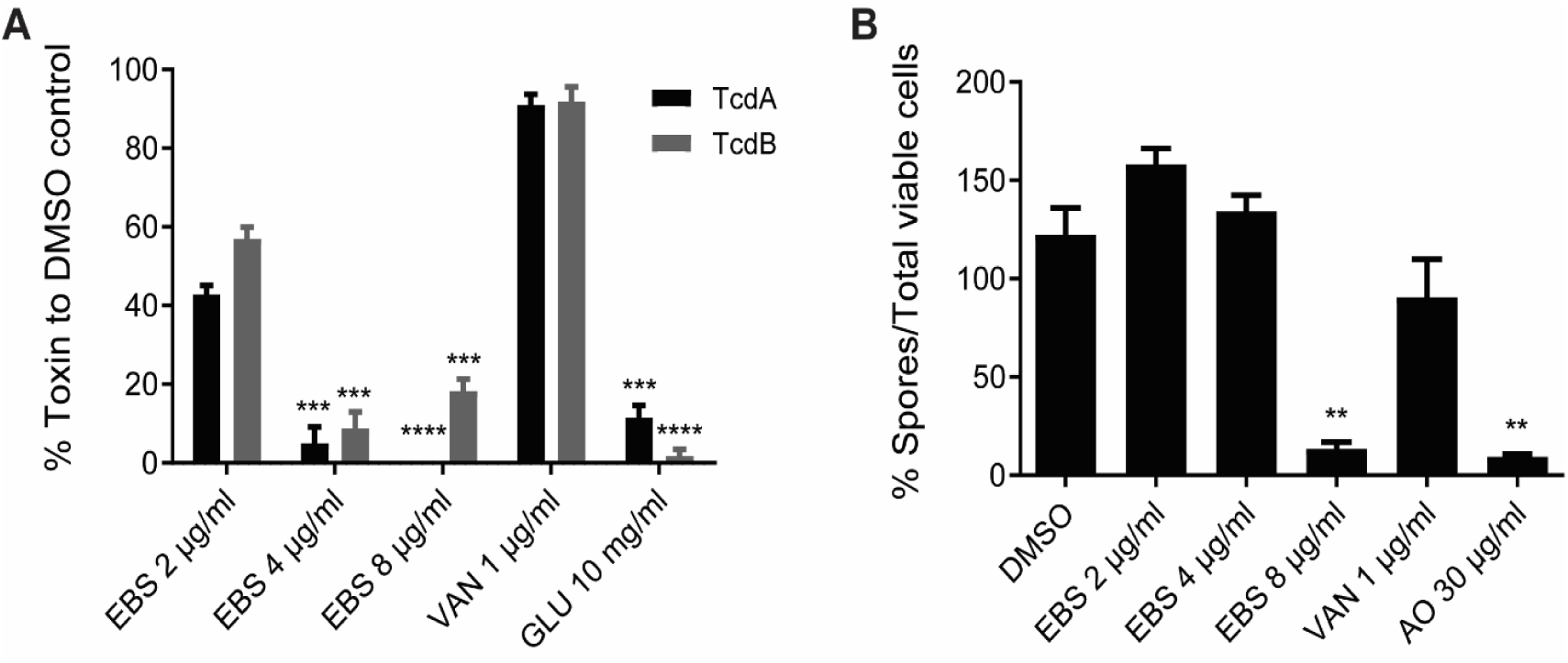
Effects of virulence by growth inhibitory concentrations of ebselen. *C. difficile* R20291 was grown to early exponential phase (OD_600_ ≈ 0.2) and treated with 2, 4 or 8 µg/ml of ebselen. **(A)** After exposure for 24 h, both TcdA (black bars) and TcdB (grey bars) were measured from culture supernatants by ELISA. Vancomycin (1 µg/ml) was a negative control and glucose (1% w/v) a positive control. Data obtained from four biological replicates were compared with respective DMSO controls. **(B)** Sporulation was analyzed after 5 days and the percentage of spores expressed per total viable population. DMSO and vancomycin (1 µg/ml) were negative controls and acridine orange (AO) at 30 µg/ml was a positive control. Error bar indicate means ± SEM; n = 3 (unpaired t-test with Welch’s correction **P<0.01, ***P<0.001 and ****P<0.0001; done using Graphpad prism version 8.4).

### Ebselen inhibits *C. difficile* spore production

Considering the significant impact of ebselen on growth of *C. difficile*, we determined whether it impacts sporulation. A mixed transcriptional response was observed for sporulation genes (**Table 3**), where mRNA for stage 0 (*spo0A*) was significantly downregulated, but stage II (*spoIIAA* and *spoIIC*) were upregulated. To directly test if ebselen hindered sporulation, we quantified spore formation in cultures exposed to ebselen for 5-days. Ebselen only inhibited sporulation at 8 µg/ml, as there was only 13.4 ± 6.8% (p = 0.003) spores in total population after treatment (**Fig. 3B**). Although, sporulation inhibition by ebselen was comparable to the positive control acridine orange (30 µg/ml), ebselen (8 µg/ml) also caused a 1 log reduction in viable cells, implying that spore reduction is coupled to cell death.

## DISCUSSION

Our findings lead to an updated model of ebselen anti-*C. difficile* properties, combining the prior report that it protects epithelial cells from TcdA and TcdB and our data showing it kills *C. difficile* cells and blocks the biogenesis of toxins and spores. Its antibacterial activity would have gone undetected save for our routine use of BHI broth, since the CLSI recommended media masks the activity of ebselen. Indeed, media for anaerobic susceptibility testing contains pyruvate and/or blood, which reduces or inactivates the activity of ebselen, respectively. There are a number of possibilities to explain this observation. Firstly, ebselen may react with thiols found in blood proteins or glutathione in blood for e.g. it is known to react with the plasma protein albumin (32). This would reduce the free fraction of unreacted ebselen for cellular activity against *C. difficile* in blood agars. Secondly, we speculate that because pyruvate is a substrate for multiple pathways, it may adjust metabolism to counter oxidative damage by ebselen. Interestingly, even in BHI media, PCR ribotype-078 strains were refractory to killing by ebselen. In theory, if the reported *in vivo* efficacy of ebselen in mice (11) arises from a combination of anti-toxin and antibacterial actions, then environmental conditions and drug resistance could hinder its activity. This may occur in CDI with intestinal bleeding and by reactions with thiol proteins such as biliary glutathione. Therefore, ebselen might need to be administered in colon specific formulations to bolster its luminal concentrations and free fraction of unmetabolized drug reaching the colon. Albeit, ebselen-resistant strains could undermine its therapeutic coverage, if antibacterial action is required for efficacy.

In determining the impact of ebselen on *C. difficile* cell physiology, we identified that it induces pathways consistent with its classification as a reactive antimicrobial. Indeed, ebselen altered cysteine metabolism, as reflected by upregulation of cystine/cysteine uptake transporters, increase cystine/cysteine content in cells and depleted the intracellular non-protein thiols. The reduction of non-protein thiols is probably due to these molecules reacting with the selenyl-sulfide bond of ebselen (33). The depletion of cellular thiols might have also been lethal to *C. difficile*, by triggering an imbalance of redox sensitive reactions. Certainly, we observed a dramatic impact on NAD^+^/NADH ratio and substantial upregulation of the proline reductase operon. Proline reductase is a selenium dependent enzyme that catalyzes reductive deamination of proline to 5-aminovalerate. It regenerates NAD^+^ from NADH (26) and this is thought to be the preferred route for NAD^+^ regeneration by *C. difficile*. PrdR, a sigma-54 dependent transcriptional regulator, activates expression of the *prd* operon and represses the *grd* operon. Upregulation of *prdR* signifies that in presence of reactive ebselen, *C. difficile* regenerated NAD^+^ through proline reduction (27). The fact that NADH concentrations were severely reduced by ebselen also supports this view, since low NADH levels favor expression of the redox dependent-transcriptional repressor *rex* (27, 34). Rex also suppresses the *grd* operon, as a less favored path for NAD^+^ regeneration in *C. difficile* (27). Currently, there is a growing impetus to better understand biological redox processes in *C. difficile* metabolism (28). We suggest that ebselen could be a useful probe to understand pathophysiological changes in response to oxidative stress.

Besides killing *C. difficile*, ebselen at its MIC (2 μg/ml) suppressed toxin production, while its inhibition of sporulation correlated with cell killing at 8 μg/ml (^1^/_2_ x its MBC). Nonetheless, both findings show that ebselen imposes secondary effects that limit *C. difficile* virulence. This is somewhat similar to observations in *Staphylococcus aureus*, where ebselen inhibited α-hemolysin production and inhibited protein, DNA, RNA and cell envelope synthesis (17). Taken together, we clearly show that ebselen’s anti-virulence properties goes beyond deactivating the CPD domain of TcdA and TcdB. In our pilot experiments, in a mouse colitis CDI model, ebselen did not rescue animals from becoming moribund when given at 10 or 100 mg/kg/day (**Fig. S2**); though animal numbers were not statistically powered. This is consistent with a recent report where ebselen did not rescue hamsters from becoming moribund due to CDI (35). In conclusion, if ebselen is to be used as a drug for CDI then chemical modification is needed to harness its polypharmacological properties and to extend its coverage to *C. difficile* ribotypes that show intrinsic antimicrobial resistance to ebselen.

## MATERIALS AND METHODS

### Bacterial strains, growth conditions and antibiotics

With the exception of *C. difficile* R20291, the various PCR-ribotypes of *C. difficile* and human gut bacteria were from Biodefense and Emerging Infectious Research Resource Repository (BEI Resources, Manassas, VA) and the American type Culture Collection (ATCC, Manassas, VA). *C. difficile* strains were grown in Brain Heart Infusion (BHI) agar or broth medium at 37°C in an anaerobic chamber (Don Whitley A35 anaerobic chamber). Other species were routinely grown in Brucella agar supplemented with 5% (v/v) defibrinated sheep blood (Hardy Diagnostics), 5 mg/L Hemin and 10 mg/L vitamin K1. The antibiotics were purchased from: Sigma (vancomycin), Acros organics (metronidazole) and Enzo life sciences (ebselen).

### Susceptibility testing

MICs were determined by the agar dilution method, as described previously (36). Briefly, serially diluted test compounds were added to molten agars of respective media and 3 µl (∼10^5^ CFU/ml) of overnight cultures were spotted onto agars using the semi-automated liquid handling benchtop pipettor (Sorenson Bioscience Inc.). After anaerobic incubation at 37°C for 24 h, the lowest concentration of compound inhibiting visible growth was recorded as the MIC. Agars used were BHI, Brucella with above supplementation and Wilkins-Chalgren. When needed BHI was also supplemented with 5% (v/v) defibrinated sheep blood, 1 g/L Na-pyruvate or hemin (5 mg/L).

### MBC determination

Cultures were grown to OD_600_nm ∼0.2 and exposed to varying concentrations of ebselen. Total viable counts were determined, at time 0 and 24 h, by plating serial dilutions of bacteria onto BHI agar plates. The MBC was defined as the lowest of compound killing 3 logs of cells from time 0.

### Transcriptome analysis

Overnight cultures of *C. difficile* R20291 were inoculated in 50 ml of fresh BHI medium and cultured until OD_600_ ≈ 0.2 (T_0_). Ebselen was added to a final concentration of 8 µg/ml and cultures incubated for a further 30 min. Cells were harvested by the addition of one volume of RNAprotect bacterial reagent (Qiagen) followed by centrifugation at 4000 x g for 10 min. Bacterial cells without ebselen treatment were prepared as a control. Cell pellets were resuspended in 700 µl of Qiazol lysis reagent (Qiagen) and lysed in a Fastprep cell disruptor at force 50 for 5 min. The total RNA was extracted using the RNeasy Mini Kit (Qiagen), according to manufacturer’s instructions. RNA Sample QC, DNase treatment, library preparations and HiSeq 2×150 Paired End sequencing were performed by GENEWIZ (South Plainfield, NJ, USA).

### Bioinformatic analysis

Raw FASTQ files were uploaded onto galaxy platform (https://usegalaxy.org/). Quality control and trimming were done using FastQC (Babraham Bioinformatics) and Trim Galore, respectively. Reads were mapped to R20291 as reference and was performed with BWA-MEM program. Counts per Read was generated with htseq-count and count matrix was generated with Join all counts program. The count matrix file was uploaded onto Degust (http://degust.erc.monash.edu/) to generate differential gene expression file using edgeR (cut-offs abs FC = 2.0 and FDR= 0.01).

### Gene expression analysis by quantitative PCR

RNA extraction was performed as indicated above. cDNA was prepared from 10 µg of total RNA using qScript cDNA supermix (Quanta Biosciences). Quantitative PCR was performed with qScript SYBR Green RT-qPCR master mix (Quanta Biosciences) using Applied Biosystems ViiA7 RT-PCR system. The C_T_ values obtained were normalized to housekeeping 16S rRNA and the results were calculated using the 2^ΔΔC^_T_ method. The primers used are shown in **Table S2**.

### Cysteine quantification

Overnight cultures of *C. difficile* R20291 were diluted 20-fold into a fresh medium and grown until OD_600_ ≈ 0.3. Compounds were added at varying concentrations and cultures incubated for 1 h. Cells were harvested by centrifugation (4000 x g for 10 min) and cell pellets resuspended in ice cold phosphate buffer. Cells were lysed in FastPrep (Qiagen) for 10 min, centrifuged at 21,100 x g for 5 min and cysteine content quantified using the Cysteine Assay Kit from Sigma-Aldrich according to manufacturer’s instructions. Cellular cysteine levels were normalized in cell lysates, by protein content determined using Pierce BCA Protein Assay Kit.

### Thiol quantification

Cell lysates obtained above were treated with 5% (w/v) trichloroacetic acid for 15 min at room temperature. Precipitated proteins were removed by centrifugation (21,100 x g for 5 min) and the pH of supernatants neutralized with 1 M Tris-base. LMW thiols were quantified using Thiol Fluorescent Detection Kit (Invitrogen) according to the manufacturer’s instructions and were normalized by protein content.

### NAD/NADH quantification

Cells were harvested as above, resuspended in PBS buffer and diluted with an equal volume of base solution (0.2N NaOH with 1% (w/v) dodecyl trimethylammonium bromide [DTAB]). After lysing in FastPrep lysates were centrifuged at 21,100 x g for 5 min. Both NAD^+^ and NADH were quantified using NAD/NADH™-GLO kit (Promega) according to manufacturer’s instructions. NAD^+^ and NADH concentrations were normalized by protein content.

### Toxin quantification

TcdA and TcdB were quantified from supernatants of 24 h cultures of R20291, using the *C. difficile* toxin A or B ELISA kit (tgcBIOMICS), according to the manufacturer’s instructions. Cultures were exposed to ebselen, vancomycin or glucose for 24 h.

### Sporulation assay

This assay was performed as described previously (37). Briefly, cultures were allowed to grow until OD_600_ ≈ 0.3 in BHI before the addition of test compounds at varying concentrations. Cultures were incubated for 5 days and sporulation evaluated as the percent of heat resistant spores per total viable count.

## ACKNOWLEDGEMENTS

This work was funded by grant R01AI132387 from the National Institute of Allergy and Infectious Diseases at the National Institutes of Health. The funders had no role in study design, data collection and interpretation of the findings, or in the writing and submission of the manuscript.

